# SnoRNA signatures in cartilage ageing and osteoarthritis

**DOI:** 10.1101/2020.04.01.019505

**Authors:** Mandy J Peffers, Alzbeta Chabronova, Panagiotis Balaskas, Yongxiang Fang, Philip Dyer, Andy Cremers, Pieter Emans, Peter Feczko, Marjolein Caron, Tim JM Welting

## Abstract

**Objectives:** Osteoarthritis (OA) presents as a change in the chondrocyte phenotype and an imbalance between anabolic and catabolic processes. Age affects its onset and progression. Small nucleolar RNAs (SnoRNAs) direct chemical modification of RNA substrates to fine-tune spliceosomal and rRNA function, accommodating changing requirements for splicing and protein synthesis during health and disease. This study was undertaken to examine how changes in snoRNAs expression may have a role in OA.

**Methods:** Articular cartilage from young, old and OA knees was used in a microarray study to identify alterations in snoRNA expression. Changes in expression of snoRNAs in OA-like conditions were studied in chondrocytes using interleukin-1 and OA synovial fluid. SNORD26 and SNORD96A knockdown and overexpression were undertaken using antisense oligonucleotides and overexpression plasmids.

**Results:** We identified panels of snoRNAs differentially expressed due to ageing (including SNORD96A, SNORD44) and OA (including SNORD26 and SNORD116) and findings were validated in an independent cohort. *In vitro* experiments using OA-like conditions affected snoRNA expression. Knockdown or overexpression of SNORD26 or SNORD96A resulted in changes in chondrogenic, hypertrophic, rRNA and osteoarthritis related gene expression.

**Conclusion:** We demonstrate that snoRNA expression changes in cartilage ageing, and OA and in OA-like conditions, and when the expression of these snoRNAs is altered this affects chondrogenic and hypertrophic gene expression. Thus, we propose an additional dimension in the molecular mechanisms underlying cartilage ageing and OA through the dysregulation of snoRNAs.

## INTRODUCTION

Osteoarthritis (OA) is the most common degenerative joint disease characterised by chondrocyte alterations and an irreversible loss of extracellular matrix (ECM) (1) leading to biomechanical failure. From a biochemical perspective, OA is characterized by uncontrolled synthesis of ECM degrading enzymes resulting in active cartilage breakdown. Chondrocytes are significant secretory cells enabling the synthesis and maintenance of the protein-rich ECM. Changes in the chondrocyte phenotype in OA are considered fundamental pathological mechanisms (2). During ageing (and joint disease), the chondrocyte’s homeostatic balance is disrupted and the rate of collagen and proteoglycan loss from the matrix may exceed deposition rate of newly synthesized molecules (reviewed (3)) resulting in increased OA risk.

To ensure continuous ECM deposition it is essential chondrocytes control the number and quality of its ribosomes. Ribosomes are cellular nanomachines equipped for conversion of genetic information encoded in mRNAs into proteins (4). Human 18S, 5.8S and 28S rRNAs contain at least 110 individual 2’O-ribose methylated and 100 pseudouridylated nucleotides (5). These post-transcriptional modifications (PTMs) fine-tune translational characteristics of the ribosome. Positioning of these modifications is undertaken by small nucleolar RNAs (snoRNAs) (6). SnoRNAs are (mainly) intron-derived non-coding RNAs of approximately 50-250 nucleotides long with an important task in the PTMs of rRNA substrates and are classified into box C/D and box H/ACA snoRNAs. Work undertaken in zebrafish suggested decreased snoRNA expression (SNORD26, SNORD44 and SNORD78) reduced the snoRNA-guided methylation of the target nucleotides and that impaired rRNA modification, at a solitary site, led to critical morphological defects and embryonic lethality. The researchers alluded that rRNA modifications play an fundamental role in vertebrate development (7). This was recently emphasized by a study showing important dynamics in rRNA ribose methylation and snoRNA expression during mouse development (8). Additionaly there are many orphan snoRNAs for which no target has been identified, however emerging evidence shows that orphan snoRNAs (and some canonical snoRNAs) fulfill non-canonical functions in alternative splicing (9), modification of other RNAs (10), a source for miRNAs (11), regulation of metabolic stress (12) and gene expression (13). Furthermore, internal 2’-O-methylation modifications (Nm) are guided by snoRNAs and these Nm sites can regulate mRNA and protein translation (14).

Atypical snoRNAs expression has been implicated in some disease processes (15) including prostate, lung and breast cancer (reviewed (16), neurodegenerative disease; Prader-Willi Syndrome (17) and viral infection (18)). We have identified aberrant expression of groups of snoRNAs in ageing cartilage (19) and diseased tendon (20). Whilst others have noted age-associated changes in snoRNAs in *C. elegans* (21). Recently through snoRNA profiling using deep-sequencing we identified differentially expressed snoRNAs relating to joint ageing and OA (22). Others have identified SNORD38 and SNORD48 as potential non-age-dependant serum biomarkers for OA progression following cruciate ligament injury (23).

Expression of snoRNAs in human articular cartilage has not been explored, neither is it known whether snoRNA expression changes in cartilage due to ageing or OA, and whether the expression of individual snoRNAs can significantly influence the chondrocyte phenotype. In this study we therefore tested the hypothesis that with age and OA there is a dysregulation of expression and function of specific snoRNAs. Our study identifies a novel set of potential OA therapeutic targets.

## METHODS

### Study design

Human cartilage was harvested from male knees at the University of Maastricht Medical Centre. Cartilage was collected at total knee arthroplasty surgery from lateral (protected (P)) and medial (unprotected (U)) femoral condyles (MUMC+ IRB; 2017-0183) or from the medial side of the lateral femoral condyle following anterior cruciate ligament repair surgery from young donors (MUMC+ IRB; 14-4-038).

### Assessment of disease severity

Old specimens came from patients with a diagnosis of OA on pre-operative knee radiographs using Kellgren and Lawrence (KL) Scoring [20]. The Outerbridge scoring system [21] was applied to U and P samples.

### Histological analysis of OA severity

Cartilage biopsies taken adjacent to those used for microarray from the protected and unprotected areas of femoral condyles were collected for histology. Biopsies were fixed in 4% paraformaldehyde and embedded in paraffin wax. 4μm longitudinal sections were cut on a microtome (Leica Biosystems, UK) and stained with Haematoxylin/Eosin (Leica Biosystems, UK) and Safranin-O/Fast Green. Sections were scored for OA severity on a scale from 1 to 11 by two independent blinded observers using a modified Mankin scoring system [22]. Inter-observer variability was calculated using Cohen’s Kappa statistics with online software (http://www.statstodo.com/CohenKappa).

### RNA extraction, microarray

RNA was extracted from cartilage [23]. Twenty-nine Affymetrix miRNA-4.0 microarrays were used. Total RNA samples were quantitated using a Nanodrop spectrophotometer (NanoDrop Technologies). 500ng of total RNA was labelled using the Affymetrix Flash-Tag Biotin HSR RNA labelling kit according to manufacturer’s instructions. Following Flash-Tag labelling the biotin-labelled samples were stored at −20°C prior to hybridisation onto Affymetrix GeneChip miRNA 4.0 for 17.5 hours at 48°C 60 rpm in an Affymetrix hybridisation oven 645. Following hybridisation the arrays were washed using Affymetrix Hybridisation wash and stain kit on the GeneChip Fluidics station 450 using fluidics script FS450_0002, and scanned using the Affymetrix GeneChip scanner 3000 7G. CEL files were generated using the Affymetrix GeneChip Command Console Software, and Expression Console software was used to QC.

### RNA isolation, poly (A) cDNA synthesis and snoRNA qRT-PCR

For validation of microarray findings an independent cohort was used. QRT-PCR of snoRNAs was performed [24]. Total RNA was isolated as described. RNA samples were polyadenylated as previously described [24]. A snoRNA-specific forward primer and a universal reverse primer (RTQ-UNIR, matched to the Tm of each snoRNA) were used for the amplification of each target (all Eurogentec, Seraing, Belgium) (Primer sequences are in Supplementary File 1). SnoRNA qRT-PCR was undertaken and the annealing temperature was optimized for each snoRNA [24]. Standard curves were used to quantify snoRNA expression and data was normalized to a validated housekeeping snoRNA. Steady-state transcript abundance of potential endogenous control genes was measured in the microarray data. Assays for SNORD63, SNORD28, SNORA28, SNORA31, U6, U2, and miR6786 were selected as potential reference genes because their expression was unaltered in the arrays.

Stability of this panel of genes was assessed with a gene stability algorithm [25] using genormPLUS (Biogazelle, Zwijnaarde, Belgium) [26] and MiR6786.

### Human chondrocyte culture studies

Human knee articular chondrocytes (HACs) were obtained following autologous cartilage repair procedures of non-OA patients (‘non-OA’) and from total knee arthroplasty of end-stage (KL grade 3-4) OA patients. Medical ethical permission was received; MEC 08-4-028. Cartilage was separated from subchondral bone and digested overnight at 37°C in collagenase type II solution (300U/ml in HEPES-buffered DMEM/F12 supplemented with antibiotics). The resulting cell suspension was rinsed with 0.9% NaCl over a 70 μm cell strainer. HAC were cultured in DMEM/F12 (Life Technologies), 10% FCS (Lonza), 1% antibiotic/antimycotic (Life Technologies) and 1% NEAA (Life Technologies) at 37°C, 5% CO_2_ and after reaching confluence cells were passaged 1:2 until passage 2. For experiments, cells were plated at 30,000 cells/cm^2^ in triplicate per donor. For non-OA (mean ±SD age years; 53±4.3, n=4) versus OA (66±2.3, n=4) human chondrocyte experiments, cells were harvested 24 hours after plating. For experiments using IL-1β (n=4) or OA synovial fluid (OA SF) (n=4), treatment started 24 hours after plating and after 24 hours of treatment cells were harvested. IL-1β (Sigma-Aldrich) was used at 10 ng/ml. OA SF (n=10) was perioperatively collected from the OA patients undergoing total knee arthroplasty and centrifuged to remove cells/debris, aliquoted and stored at −80°C. For experiments, SF from 10 patients was pooled and applied on cells in a 20% (v/v) concentration. Details of all donors are in Supplementary file 2. The expression of SNORD116, SNORD96A, SNORD26, SNORD33, SNORD44, SNORD95 and SNORD98 were measured in HAC with qRT-PCR using methods described previously. A panel of chondrocyte phenotype genes was measured in these samples; SOX9, ACAN, COL2A1, RUNX2, COL10A1, MMP13 and COX2. To determine the effect of oxygen tension (5% or 20%) and serum (serum-free or 10% foetal calf serum (FCS)) on snoRNA gene expression non-OA HACs were plated at 30,000 cells/cm^2^ in triplicate per donor (passage 1, n=3 males; mean±standard deviation age, 25±20), using methods above. RNA was harvested 48 hours after plating. The extent of chondrocyte dedifferentiation due to passage on snoRNA expression was assessed using equine articular chondrocytes isolated from grossly normal metacarpophalangeal cartilage (n=5; mean±standard deviation age, 4±1). Cells were cultured as described above in 20% oxygen in 10% FCS and passaged at 80% confluence from P0 to P3. Selected snoRNAs DE in the microarray were measured along with chondrogenic, and hypertrophic genes.

### Topographical changes in snoRNA expression

To assess topographical changes in snoRNA gene expression we utilised cartilage from high and low load areas of the equine metacarpophalangeal joint. Donor RNA (n=5;age mean±standard deviation 7.6±0.9 years old) was extracted from cartilage of grossly normal medial and lateral condylar regions of the joint representing high and low load areas (24). Selected snoRNAs DE in the microarray were measured along with chondrogenic, and hypertrophic genes.

### Microarray data analysis

Expression Console software was utilised for array quality control. CEL files were generated with the Affymetrix GeneChip Command Console Software. The snoRNA expression data measured using Affymetrix miRNA 4.0 arrays were preprocessed using Affymetrix Expression Console with optioned method RMA [27] for normalisation. The further statistical analyses were carried out on the 1996 snoRNA and scaRNA probe sets for *Homo sapiens* extracted from all probes and used to determine the detected and differentially expressed (DE) snoRNAs. In each test the p-value of each sample was combined using Fisher’s combined p-value methods. The expressions were dereplicated to transcript level by averaging replicated probes. The p-value associated with the presence of dereplicated expression was assigned by combining replicated probes using Fisher’s combined p test. The DE analyses on contrasting sample conditions were performed through linear models using limma package [28] in R. The significance of log fold change (logFC) values for snoRNAs were evaluated using t-tests, the p-values associated with logFC values were adjusted for multiple testing using the False Discovery Rate (FDR) [29]. Significantly DE was defined as those with an FDR-adjusted P-value < 5%.

### SNORD26 and SNORD96A knockdown using antisense oligonucleotides (ASOs)

HACs from six non-OA donors (Supplementary File 2) seeded individually at 30,000 cells/cm^2^ in triplicate were transfected (HiPerfect; Qiagen, Manchester, UK) with 100 nM ASO (Eurogentec, Belgium) targeting SNORD26 or SNORD96A. A scrambled SNORD26 or SNORD96A ASO were used as controls (Supplementary File 1). Cells were harvested after 24 and 48 hours for RNA (25). Expression of SNORD26, SNORD96A and a panel of chondrogenic and hypertrophic markers were measured relative to cyclophilin (26).

Alkaline phosphatase (ALP) was determined following lysis of a parallel well per donor, condition and time point using an ALP activity assay using p-nitrophenyl phosphate (pNPP) as a phosphatase substrate. Data was corrected for DNA content. Prostaglandin E2 (PGE2) secretion was determined in culture medium at 24 and 48 hours using an ELISA (Cayman Chemicals, USA) (26).

### SNORD26 and SNORD96A overexpression

The sequences of human SNORD26 and SNORD96A, flanked by MluI and XhoI restriction sites (Supplementary File 1), were synthesized and cloned into pUC57 by GeneCust (Ellange, Luxembourg). SnoRNA sequences were then sub-cloned into the pCGL-1 plasmid (kind gift of Prof. Tamás Kiss (27)) using MluI and XhoI restriction sites in the second intron of β-globin in pCGL-1. Primary HAC (pool of 5 donors,) (Supplementary File 2) seeded at 40,000 cells/cm2 were transfected with 2 μg pCGL-1-SNORD26, pCGL-1-SNORD96A or empty pCGL-1 plasmid using Fugene (Promega, Madison, WI, USA). Cells were harvested 24- and 48-hours post-transfection for RNA isolation (TRIzol; Invitrogen). Gene expression data were normalized to cyclophilin.

### Statistical analysis

Statistical evaluation of KL and histological scoring, and topographical gene expression results were undertaken using a Mann-Whitney Test. Other data were tested for normality prior to further analysis using Student’s t-test to compare between two samples, or one-way ANOVA with *post hoc* Tukey’s test to compare between multiple samples. Statistical analysis was performed in GraphPad Prism version 8.03 (GraphPad, La Jolla California USA, www.graphpad.com); p-values are indicated.

### Data availability

Data are deposited in NCBI’s Gene Expression Omnibus (GEO Series accession number E-MTAB-5716).

## RESULTS

### OA grading and histological characterisation of articular cartilage

Young cartilage had no gross signs of OA. To confirm the OA protected (P) and OA unprotected (U) nature of cartilage, KL scoring revealed a significant increase in KL score in P (mean±standard deviation) (1.9±0.8) compared to U (3.0±0.8) with p=0.03, and in Outerbridge Score in P (1.2±0.9) compared to U (2.3±1.2) with p=0.0016. Modified Mankin scores for P; 2.6±1.3 and U; 4.8±2.6, p=0.04 (data not shown). Cohen’s Kappa statistic was 0.6 indicating a strong agreement between observers. As a result, potential snoRNAs expression differences between the P and U samples may be attributed to OA severity scores. Supplementary File 3 summarises gross score, Modified Mankin, Outerbridge, and KL scores of donors.

### Microarray snoRNA analysis overview

RNA was extracted and used in microarray studies from ten donors for P and U and nine for Y. Data quality assessment of the data was good and consistent for all arrays. The outcomes of variation assessment are visualised in Figures 1A and 1B. Sample clustering is demonstrated by the correlation coefficient matrix heatmap (Figure 1A). A Principal Component Analysis (PCA) plot confirmed that young samples were separated from old (P and U) samples. Samples from the P group clustered into two sub-populations; designated P1 and P2 (Figure 1B). P1 contained four donors (samples 2, 4, 6, 8) with age (mean±standard deviation) 55.5±3.5 and P2 contained six donors (samples 10, 12, 14, 16, 18, 20) age 67.3±4.6, p=0.01. There was no difference in the KL scores or the Modified Mankin scores between P1 and P2 (data not shown). Given no other available clinical data for the donors we hypothesize that age accounts for snoRNAs expression differences between P1 and P2. The old OA (U) samples were scattered between U1, U2 and U3 clusters. Samples 5, 11 and 19 (U1) were most similar to samples from the young group. U2 consisting of samples 9 and 13 and 17 were similar to P2 (Figure 1B). U3 contained samples 1, 3, 13, 15. There was no statistical difference in age, Modified Mankin scores or KL scores between U1, U2 or U3 (data not shown).

**Figure 1.**
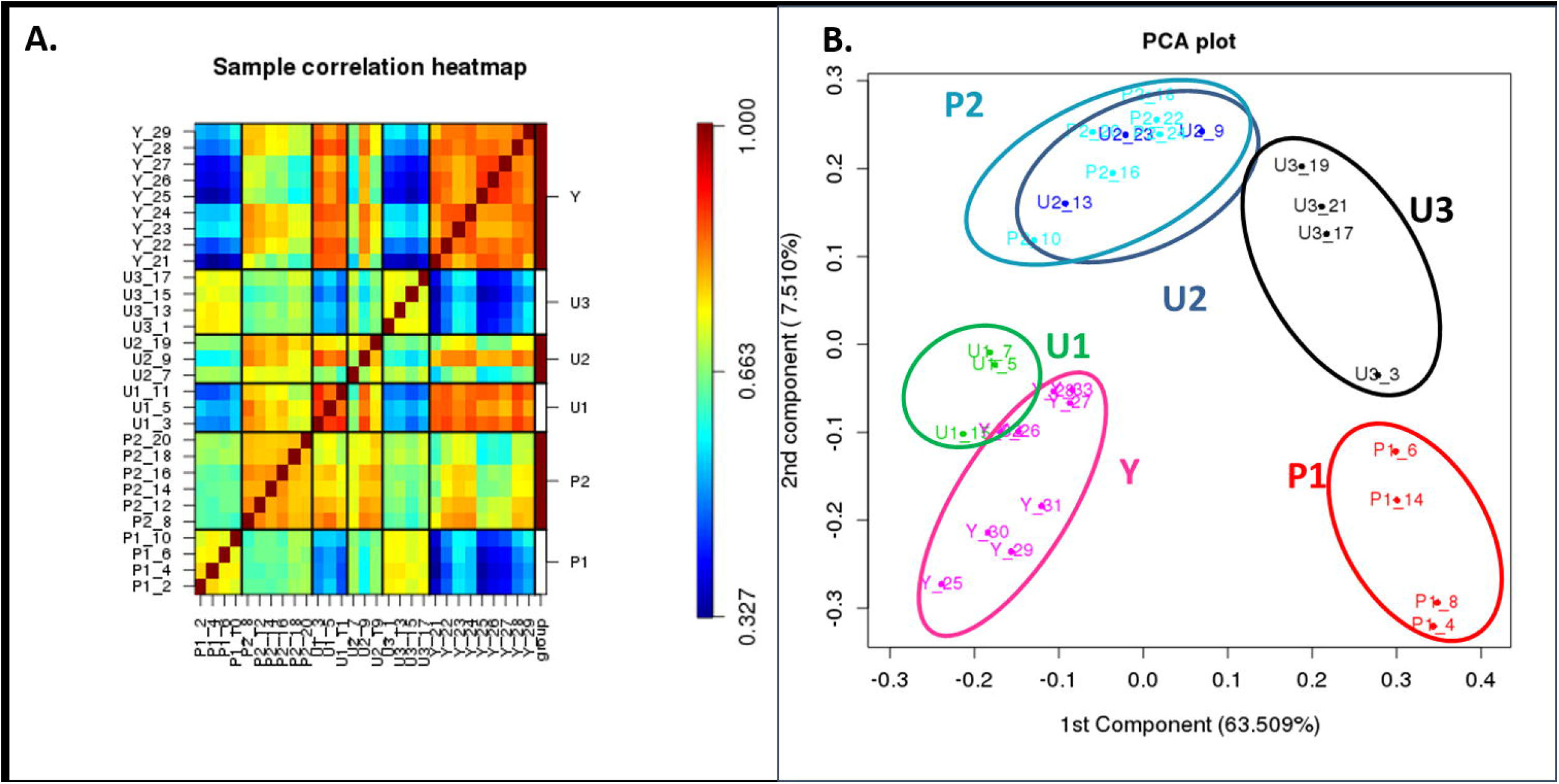
Variation of data between the expressions for 29 microarray samples. **A.** The heat map of hierarchical clusters of correlations among samples. Pearson’s correlation coefficients were computed using logarithm transformed snoRNAs expression data from all snoRNA probes detected. **B**. A 2-D PCA plot of the first and second components from PCA of logarithm-transformed snoRNA expression data. PCA plot was generated after centring each gene expression to zero in order to assess the effects of the factors. Samples represented as three groups; young (Y), protected (P) and unprotected (U).

### Differential expression of snoRNAs

Of the 1996 human snoRNA and scaRNA probes represented on the Affymetrix GeneChip miRNA-4.0 microarray, 297, 378 and 368 were detected above background in Y, P and U, respectively (Supplementary File 4).

We identified panels of snoRNAs DE. As we were interested in DE snoRNAs in ageing and OA, and the old (P) samples were in two distinct groups (P1 and P2), we made the following contrasts; for ageing changes; Y versus P1, Y versus P2, and P1 versus P2; and for OA-related changes P1 versus U and P2 versus U. The number of DE snoRNAs with FDR<0.05 are in Table 1. SnoRNAs DE in both Y versus P1 and Y versus P2 are in Supplementary File 5.

**Table 1.** Number of differentially expressed snoRNAs between contrasts

| Contrast definition | young versus old OA protected | young versus old OA protected | old OA protected versus old OA unprotected | old OA unprotected versus old OA protected | age |
| --- | --- | --- | --- | --- | --- |
| Contrast | Y vs P1 | Y vs P2 | U vs P1 | U vs P2 | P1 vs P2 |
| No. DE | **126** | **39** | **39** | **2** | 52 |
| No. Increased | **17** | **35** | **33** | **2** | 4 |
| No. Decreased | **109** | **4** | **6** | **0** | 48 |

Abbreviations- Y; young normal, P; protected (old OA protected), U; unprotected (old OA unprotected)

### Validation of differentially expressed snoRNAs

Microarray data was validated using an independent cohort. Seven selected DE snoRNAs were assessed, based on the level of DE and following a literature search of disease-related snoRNAs. Validation performed in a set of eight Y, seven P and seven matched U samples showed comparable effects and direction of DE (Figures 2A and 2B). Figure 2B demonstrates the comparative features within the microarray and qRT-PCR results. Topographical area did not affect selected snoRNA expression (Supplementary File 6). Together these data show that in HAC snoRNAs are DE as a function of ageing and OA.

**Figure 2.**
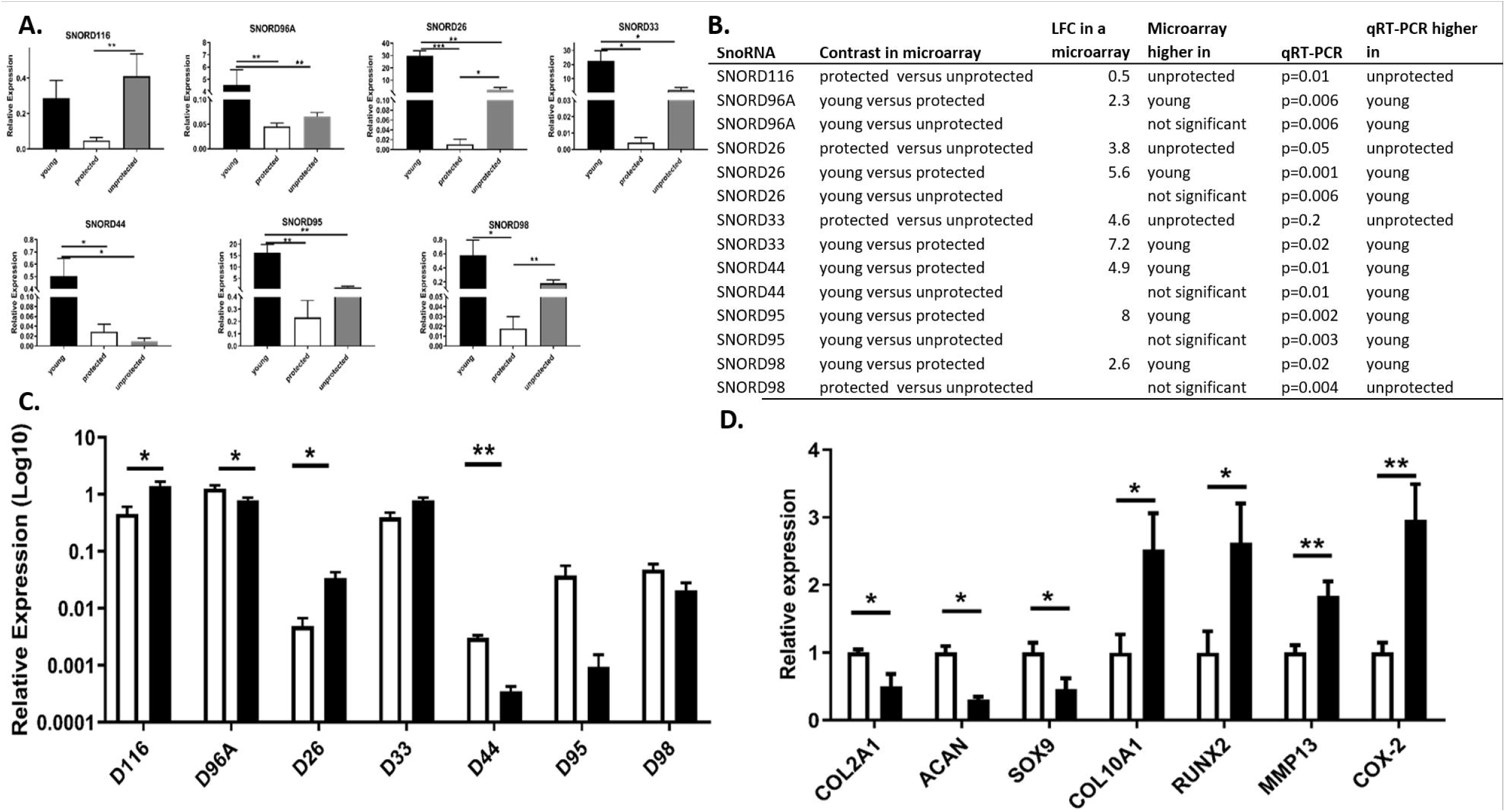
SnoRNA expression in cartilage and OA chondrocytes. **A**. Selected snoRNAs DE in the microarray were validated using qRT-PCR. Gene expression patterns of SNORD95, SNORD96A, SNORD26, SNORD98, SNORD33, and SNORD44 were validated in Y versus P cartilage. SNORD116 and SNORD26 and SNORD98 were validated in P versus U. SNORD96A, SNORD26, SNORD33, SNORD95 and SNORD44 were differentially expressed between Y and U cartilage. Only the latter was DE in the microarray analysis. **B**. The table shows a comparison between the microarray and qRT-PCR results for selected snoRNAs. All microarray results had an FDR<0.05. **C**. The gene expression of selected snoRNAs DE in the microarray were determined in non-OA (n=4) and OA (n=4) human articular chondrocytes. SnoRNA expression; SNORD116, SNORD26, SNORD33, SNORD95 were increased in ‘protected’ compared to ‘unprotected’ cartilage. SNORD96A, SNORD44 and SNORD98 were reduced in ageing cartilage. **D**. The chondrocyte phenotype differences between these donors were assessed using a panel of chondrocyte phenotypic genes. Gene expression changes were measured using 2^-ΔCT expression relative to miR6786 (snoRNAs) or cyclophilin (protein coding genes). A logarithmic scale was used to compress the data were appropriate. White bars represent non-OA and black OA chondrocytes. Data represents the mean + standard error of mean, p-values are indicated. P-values indicated as follows; p<0.05 *, p<0.01 **.

### Selected snoRNAs expression in OA chondrocytes and following IL-1β or synovial fluid treatment of non-OA chondrocytes

We measured seven snoRNAs DE in the microarray in isolated non-OA and OA HACs and following treatment of a pool of non-OA HACs with IL-1β or OA synovial fluid. There was a significant increase in the age of OA compared to non-OA chondrocytes (p=0.01). We detected a significant increase in expression of SNORD116 and SNORD26 in OA chondrocytes (increased in P versus U in the microarray). SNORD96A and SNORD44 expression was reduced in OA chondrocytes (reduced in Y versus P in the microarray). No significant change in expression was demonstrated in SNORD98 or SNORD33 (reduced in Y versus P in the array) (Figure 2C). There was an increase in expression of SNORD116, SNORD26, SNORD33 and SNORD98 following IL-1β treatment (Figure 3A). Finally, we assessed the effect of OA SF on a pool of non-OA chondrocytes. There was an increase in expression of SNORD116, SNORD96A, SNORD26, SNORD33, SNORD95 and SNORD98 (Figure 3B). Upon exposure to IL-1β or OA SF there was reduced expression of chondrogenic genes (COL2A1, ACAN, SOX9) and increase in hypertrophic genes (COL10, RUNX2, MMP13) (Supplementary File 7).

**Figure 3.**
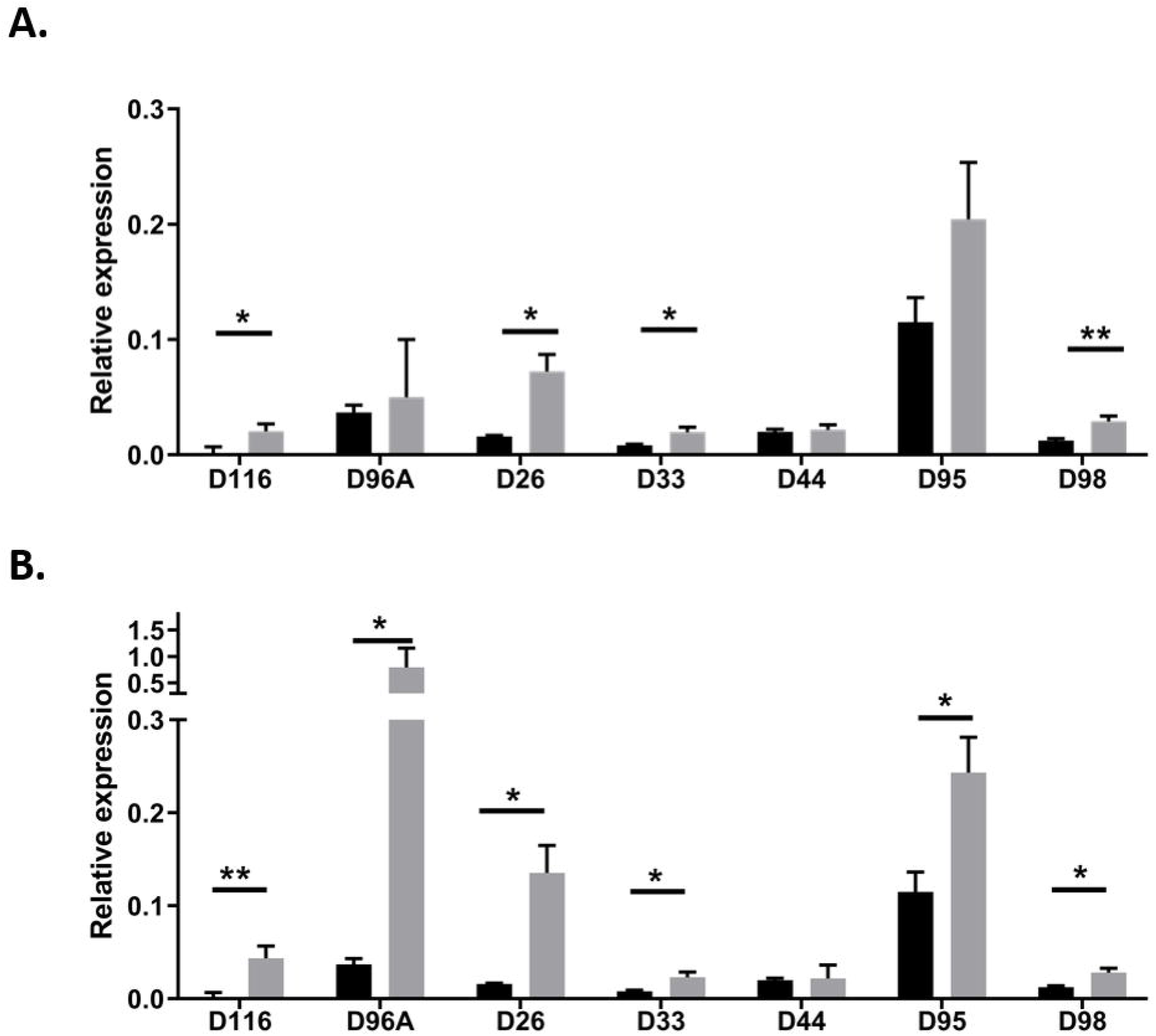
Selected snoRNA expression in OA-like conditions. **A.** SnoRNA expression following IL-1β treatment of non-OA HAC pool of n=4 donors. Gene expression of SNORD116, SNORA26, SNORD96A, SNORD33, SNORD44, SNORD66, SNORD95, SNORD98 were was measured using qRT-PCR following treatment of non-OA HACs with IL-1β (10 ng/ml) for 24 hours. **B.** SnoRNA expression following treatment of non-OA HAC pool of n=4 donors with 20% OA synovial fluid (SF) (derived from a pool of ten donors) for 24 hours. Gene expression changes were measured using 2^-ΔCT expression relative to miR6786. Data represents the mean + standard error mean, P-values indicated as follows; p<0.05 *, p<0.01 **. Within the graphs black represents control and grey treatments.

### SNORD26 and SNORD96A knockdown

To determine whether interference with the expression of a single snoRNA induces cellular changes relevant for the chondrocyte phenotype, we reduced expression of SNORD26 or SNORD96A using ASOs in HACs. A significant reduction in SNORD26 (24 hour; 60%, 48 hour; 61%), and SNORD96A (24 hour; 72%, 48 hour; 69%) was confirmed following target-specific ASOs transfection. Following both SNORD26 and SNORD96A knockdown at 24 and 48 hours there were significant alterations in expression of rRNAs, chondrogenic, hypertrophic and OA-related genes (Figure 4A-D). To functionally confirm hypertrophy-related gene expression changes (28) at the protein level following SNORD26 and SNORD96A knockdown, we measured PGE_2_ in culture supernatants (as a confirmation of COX2 expression) and ALP activity in cell extracts (as a confirmation of ALPL expression). We found an increase in COX2 expression was functionally accompanied by increased PGE2 in supernatants and increased ALPL expression accompanied the same alterations in ALP enzymatic activity (Figure 4E-H).

**Figure 4.**
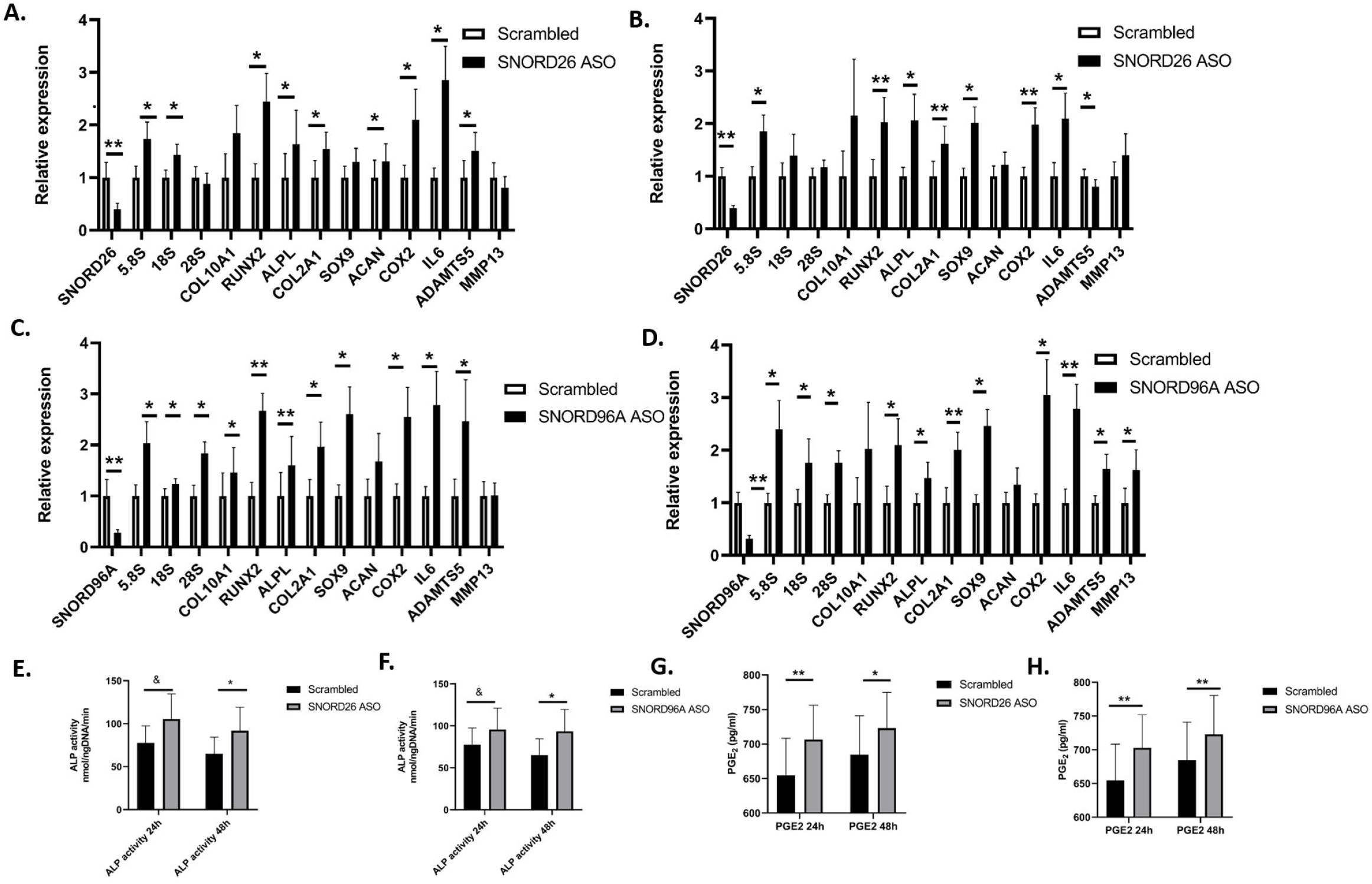
Antisense oligonucleotide-mediated knockdown of SNORD26 and SNORD96A. HACs (n=6) were transfected with either 100 nM SNORD26 or SNORD96A anti-sense oligonucleotides (ASO, black bars) or with scrambled version (white bars) of this ASO. **A**. 24-hour SNORD26, **B.** 48-hour SNORD26, **C**. 24-hour SNORD96A, **D.** 48-hour SNORD96A. SnoRNA, rRNAs, hypertrophic, chondrogenic and OA genes were determined by qRT-PCR. Alkaline phosphates activity was measured in cell lysates of parallel wells following **E**. SNORD26 knockdown for 24 and 48 hours; **F.** SNORD96A knockdown for 24 and 48 hours. PGE2 protein was measured in media following **G.** SNORD26 knockdown for 24 and 48 hours; **H.** SNORD96A knockdown for 24 and 48 hours. Statistical evaluation was undertaken using an independent samples t-test relative to the corresponding control condition using GraphPad Prism. The p-values are indicated; & p<0.06, * p<0.05, ** p<0.01. Data represents the mean value of 6 technical replicates and error bars represent standard error mean.

### SNORD26 and SNORD96A overexpression

Reciprocally, we increased expression of SNORD26 and SNORD96A individually into primary chondrocytes by transfection of engineered snoRNA mini-genes and measured gene expression 24 hours and 48 hours later. We produced a profound overexpression of each snoRNA with similar trends for protein coding gene expression at each time point. However, the most pronounced effect was evident at 48 hours. At 24 hours there were alterations in the OA chondrocyte-relevant genes COX2 and IL6 (SNORD26) and COX2 and MMP13 (SNORD96A). At 48 hours after transfection of the snoRNA mini-genes, for both SNORD26 and SNORD96A we identified significant changes in expression of chondrogenic, hypertrophic and OA-relevant genes and of 18S and 28S rRNAs for SNORD26 (Figure 5A-D). Together with our knockdown data we demonstrated that alterations in the expression of SNORD26 and SNORD96A in primary HAC profoundly changes the chondrocyte’s phenotype and rRNA expression.

**Figure 5.**
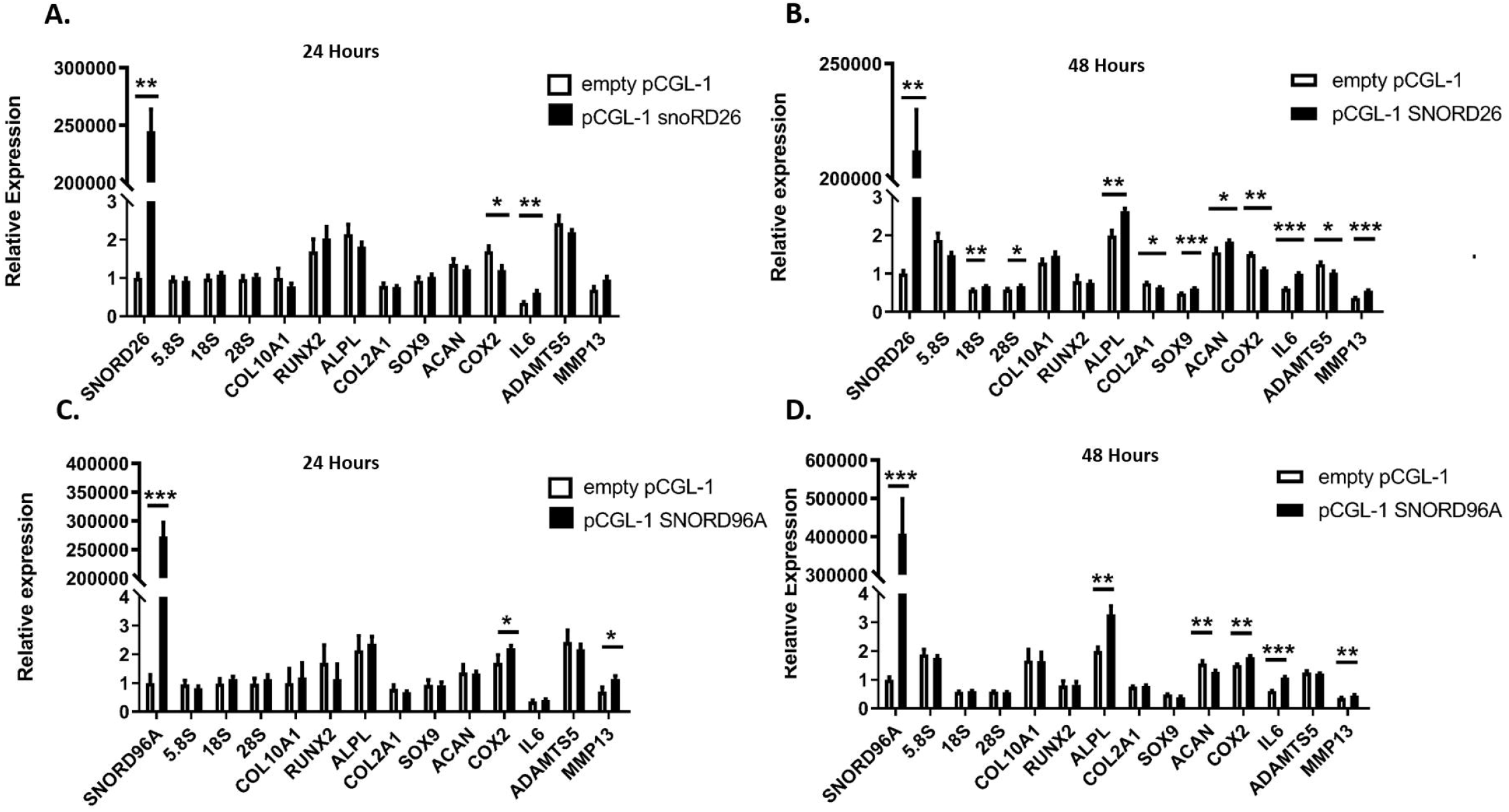
Overexpression of SNORD26 and SNORD96A. Primary HACs (n=5 pool) transfected with SNORD26 and SNORD96A mini genes (black bars) or empty vector (white bars). **A.** 24-hour SNORD26, **B.** 48-hour SNORD26, **C.** 24-hour SNORD96A, **D.** 48-hour SNORD96A. SnoRNA, rRNAs, hypertrophic, chondrogenic and OA genes were determined by RT-qPCR. Data is presented as relative to the control condition (empty vector) and for statistical evaluation an independent samples t-test was performed relative to the corresponding control condition using GraphPad Prism. The p-values are indicated; * p<0.05, ** p<0.01, ***p<0.001. Data represents the mean value and error bars represent standard error mean.

## DISCUSSION

There is increasing evidence that snoRNAs have important roles in cellular development, homeostasis and disease. Indeed we previously demonstrated novel snoRNAs features in joint ageing and OA in the mouse (22), and cartilage ageing in the horse (19). Furthermore, we have reported that the expression of the non-canonical snoRNA RMRP is regulated during chondrogenic differentiation and regulates chondrocyte hypertrophy (29). The snoRNA field is a *terra incognita* regarding knowledge on the functional implications of individual snoRNAs in cell biological processes, and this is a novel area of OA research. In OA the balance between cartilage ECM anabolism and catabolism is disrupted and important alterations in the chondrocyte phenotype are evident (2, 30). The maintenance of the ECM demands a sufficient number of functional ribosomes to translate cartilage ECM-related mRNAs into proteins and ribosome functionality may also adapt to OA chondrocyte phenotypic changes. SnoRNAs are essential for ribosome function by PTMs of rRNAs, which critically supports rRNA stability, inter-molecular interactions between rRNA and ribosomal proteins, and are essential for the ribosome’s peptidyl transferase activity and mRNA decoding activity (20, 28). Here we identified for the first time that canonical snoRNA expression changes during cartilage ageing and OA and propose an additional dimension in the molecular mechanisms underlying cartilage ageing and OA. These changes in cartilage snoRNA expression appeared to be specific of/for ageing and OA, since we demonstrated there was little effect on chondrocyte snoRNA expression of hypoxia, serum, passage number and location of cartilage within the joint (Supplementary Files 6, 8, 9).

Our study identified that SNORD26 was increased in OA and SNORD44 and SNORD78 were reduced in ageing. Previously it has been reported that reduction of snord26, snord44 and snord78 leads to morphological abnormalities and lethality in zebra fish development (7). Also the expression of snord78 has been reported to depend on mouse development, with concomitant changes in rRNA target methylation and potential consequences for ribosome specialisation (8). Additionally, further evidence of the biological relevance of DE snoRNAs is evident for SNORA40 (DE in ageing), targeting helix 27 in 18S rRNA and controlling the ribosome’s decoding function (31). Helix 68 (part of the ribosome’s E site in 28S rRNA) has an important role in protein translation via stabilisation of the peptidyl transferase centre (32). In our study SNORD36 (DE in ageing and OA), SNORA31, SNORA27, SNORD87 and SNORD88 (all DE in ageing) all target helix 68 modifications. Although most of the rRNA work in relation to PTMs and ribosomal fidelity has been undertaken in yeast, rRNA modification patterns are largely maintained throughout evolution, enabling projection from yeast to human (33). Thus, we hypothesize that the DE of canonical snoRNAs in concert alters ribosome functionality, relevant for maintaining cartilage tissue homeostasis and a healthy chondrocyte phenotype. We were therefore intrigued by the fact that OA-relevant extracellular environments like IL-1β and OA SF altered snoRNA expression and we speculate that age- and OA-associated DE snoRNAs impact ribosomal function and thereby the translational control (34) relevant for cartilage homeostasis.

We measured the expression of selected snoRNAs from the microarray data in an independent cohort and in cultured non-OA and OA chondrocytes. From these selected snoRNAs increased expression in chondrocytes isolated from OA cartilage was evident for SNORD26, which is in concert with increased expression found in OA cartilage. The expression of SNORD96A was heavily reduced due to age in cartilage, and slightly reduced in isolated OA chondrocytes. These findings suggest that differential expression of SNORD26 is predominantly OA-related, whereas expression of SNORD96A is predominately age-related. We investigated the relevance of these two snoRNA in detail. The knockdown of SNORD26 and SNORD96A appeared to have similar gross transcriptional effects, predominately increasing OA, chondrogenic and hypertrophic gene expression and causing an overall upregulation of rRNA expression. SNORD96A knockdown is a relevant condition for the observed age-related reduction in cartilage. The apparent overall deregulation of chondrocyte gene expression following its knockdown indicates an age-related central function of SNORD96A in chondrocyte homeostasis. The overexpression of SNORD26 or SNORD96A led to more distinct snoRNA-specific differences. For both SNORD26 and SNORD96A the effect of overexpression on the mRNAs measured was most pronounced at 48 hours after transfection, expected due to the time required for the β-globin gene to be transcribed, splice and ‘release’ the snoRNA. SNORD26 overexpression is a relevant condition for the here observed OA-related induction of cartilage SNORD26. Indeed its overexpression induced the expression of many OA-related genes, with an apparent uncoupling of expression of chondrogenic genes (increase in SOX9 and ACAN, but a reduction in COL2A1). In contrast to SNORD26, SNORD96A overexpression was less disruptive for chondrocyte transcriptional homeostasis, with mainly ALPL responding and a weak effect on COX2, IL6 and MMP13.

Non-canonical and orphan snoRNAs have no (predicted) function in the PTMs of rRNAs, have a variety of non-canonical functions. SNORD32A, SNORD33 and SNORD35A have non-canonical roles as critical promoters of metabolic stress. A loss of these snoRNAs causes resistance to lipotoxic and oxidative stress *in vitro* and prevents prorogation of oxidative stress *in vivo* (11). These snoRNAs shuttle to the cytoplasm and trigger cell death in response to oxidative stress. In our study we found an increase in OA and/or ageing of SNORD33 and SNORD35A and SNORD35A. Additionally there was increased expression of SNORD33 following OA SF treatment of HACs. Oxidative stress in cartilage ageing and OA development has been reported (35). We speculate that the OA and/or ageing-dependent increased expression of these oxidative stress-related snoRNAs is a representation of oxidative stress pathway activity. We identified SNORD116 as another DE non-canonical snoRNA in OA. SNORD116 also responded to IL-1β and OA synovial fluid treatment. In concert with the upregulation of SNORD116 in OA cartilage we previously reported the upregulation of SNORD116 in the mouse DMM model (22). Microdeletions of the SNORD116 cluster is evident in Prader-Willi Syndrome (PWS), a disease leading to developmental delay and genetic obesity (36). SNORD116 lacks any significant complementarity with rRNA targets and has been mainly implicated in alternative splicing of specific target genes (13). The significance of the DE of SNORD116 in OA cartilage remains to be determined, but an inflammation-related regulation of alternative splicing in chondrocytes is expected to be an interesting avenue for further investigation (37).

In this study we wished to determine snoRNA expression in cartilage ageing and OA. We were unable to source normal cartilage from old joints without any gross signs of OA within the entire joint. Thus, a limitation of this study is that we were unable to collect truly ‘normal’ old cartilage. Instead we used matched cartilage from the lesser affected lateral condyles of knees removed following TKA, as previously suggested [28]. Indeed, there was an increase in KL, Outerbridge and Modified Mankin’ scoring in OA compared to old. Our data also demonstrates that snoRNA expression in chondrocytes residing in minimal OA cartilage have a distinct pattern of gene expression compared to those residing in advanced OA cartilage irrespective of gender or age.

This study comprehensively determined snoRNA signatures in ageing and OA cartilage.

Given canonical but also emerging non-canonical functions of snoRNAs in homeostasis and disease, snoRNAs provide a plethora of unexplored molecular routes in chondrocyte biology. This warrants further in-depth studies addressing the cellular functions of specific snoRNAs in chondrocyte pathobiology and explore their potential as novel molecules to target OA treatment.

## ACKNOWLEDGMENTS

We thank Mandy Steinbusch for her assistance with laboratory work. Mandy Peffers is funded through a Wellcome Trust Intermediate Clinical Fellowship (107471/Z/15/Z). Tim Welting is funded through Dutch Arthritis Society grants LLP14 and 17-2-401, NWO-DFG grant 82-304 and Stichting de Weijerhorst. Some of the work from this manuscript has previously been presented at Osteoarthritis Research Society International 2018 (Osteoarthritis and Cartilage 26: S164. Apr 2018).

