## Supplementary 1 for "SnoRNA signatures in cartilage ageing and osteoarthritis"

Supplementary File 1.

A qRT-PCR primers sequences used

| **Species** | **Primer Name** | **Forward** | **Reverse** |
| --- | --- | --- | --- |
| Equus caballus | gapdh | GCA-TCG-TGG-AGG-GAC-TCA | GCC-ACA-TCT-TCC-CAG-AGG |
| Equus caballus | 5.8S rRNA | GCG-TTC-CTT-GGC-TGG-TTC-A | AGT-GCG-AAG-TGT-CCA-TGA-CC |
| Equus caballus | 18S rRNA | CGG-ACA-GGA-TTG-ACA-GAT-TGA-TA | TGC-CAG-AGT-CTC-GTT-CGT-TA |
| Equus caballus | 28S rRNA | CGG-GTA-AAC-GGC-GGG-ACT-AAC | TAG-GTA-GGG-ACA-GTG-GGA-ATC-TCG |
| Equus caballus | sox9 | CTC-TGA-ATG-TCA-AAG-TGA-AGA-AAG-T | GAC-GAG-GCA-TTT-GGC-TAC-C |
| Equus caballus | aggrecan | GAG-GAG-CAG-GAG-TTT-GTC-AAC-A | CCC-TTC-GAT-GGT-CCT-GTC-AT |
| Equus caballus | col2a1 | TCA-AGT-CCC-TCA-ACA-ACC-AGA-TC | GTC-AAT-CCA-GTA-GTC-TCC-GCT-CTT |
| Equus caballus | col1a1 | CAT-GTT-CAG-CTT-TGT-GGA-CCT | TGA-CTG-CTG-GGA-TGT-CTT-CTT |
| Equus caballus | mmp13 | CTG-GAG-CTG-GGC-ACC-TAC-TG | ATT-TGC-CTG-AGT-CAT-TAT-GAA-CAA-GAT |
| Equus caballus | runx2 | CAGACCAGCAGCACTCCATA | CAGCGTC |
| Homo sapiens | GAPDH | ATG-GGG-AAG-GTG-AAG-GTC-C | AACACCATCATTC |
| Homo sapiens | 5.8S rRNA | CAC-TCG-GCT-CGT-GCG-TCG-AT | CGC-TCA-GAC-AGG-CGT-AGC-CC |
| Homo sapiens | 18S rRNA | CGG-ACC-AGA-GCG-AAA-GCA | ACC-TCC-GAC-TTT-CGT-TCT-TGA-TT |
| Homo sapiens | 28S rRNA | GCC-ATG-GTA-ATC-CTG-CTC-AGT-AC | GCT-CCT-CAG-CCA-AGC-ACA-TAC |
| Homo sapiens | COL1A1 | GTT-CAG-CTT-TGT-GGA-CCT-CCG | GAT-TGG-TGG-GAT-GTC-TTC-GTC-T |
| Homo sapiens | COL2A1 | TGG-GTG-TTC-TAT-TTA-TTT-ATT-GTC-TTC-CT | GCG-TTG-GAC-TCA-CAC-CAG-TTA-GT |
| Homo sapiens | COL10 | ATG-ATG-AAT-ACA-CCA-AAG-GCT-ACC-T | ACG-CAC-ACC-TGG-TCA-TTT-TCT-G |
| Homo sapiens | aggrecan | TCG-AGG-ACA-GCG-AGG-CC | CG-AGG-GTG-TAG-CGT-GTA-GAG-A |
| Homo sapiens | SOX9 | GAG-CAG-ACG-CAC-ATC-TC | CCT-GGG-ATT-GCC-CCG-A |
| Homo sapiens | ADAMTS5 | GTG-GCT-CAC-GAA-ATC-GGA-CAT | GCG-CTT-ATC-TTC-TGT-GGA-ACC-A |
| Homo sapiens | IL6 | CGA-GAA-AAC-AAC-CTG-AAC-CTT | ACC-TCA-AAC-TCC-AAA-AGA-CCA |
| Homo sapiens | cox2 | ACC-AAC-ATG-ATG-TTT-GCA-TTC-TTT | GGT-CCC-CGC-TTA-AGA-TCT-GTC-T |
| Homo sapiens | MMP13 | CTT-CAC-GAT-GGC-ATT-GCT-GAC | CGC-CAT-GCT-CCT-TAA-TTC-CA |
| Homo sapiens | ALPL | CCG-TGG-CAA-CTC-TAT-CTT-TGG | CAG-GCC-CAT-TGC-CAT-ACA-G |
| Homo sapiens | CYCLOPHILIN | TTC-CTC-CTT-TCA-CAG-AAT-TAT-TCC-A | CCG-CCA-GTG-CCA-TTA-TGG |
| Homo sapiens | RUNX2 | TGA-TGA-CAC-TGC-CAC-CTC-TGA | GCA-CCT-GCC-TGG-CTC-TTC-T |

**SNORNA forward primer sequences**

| **Species** | **Primer Name** | **Forward** |
| --- | --- | --- |
| Homo sapiens | snord63 | GAA-AGA-ACG-TGT-GGA-AAA-CTA-ATG |
| Homo sapiens | U6 | GAT-GAC-ACG-CAA-ATT-CGT-GAA-GCG-TTC |
| Homo sapiens | U2 | TGG-TAT-TGC-AGT-ACC-TCC-AGG-AAC-G |
| Homo sapiens | 116_1 | ACC-AAA-CCA-CTT-CTG-TGA-GCT-G |
| Homo sapiens | 116_30 | ACC-AAA-CCA-CTT-CTG-TGA-GCT-G |
| Homo sapiens | snord13 (U13) | AAC-CTT-GTT-ACG-ACG-TGG-GCA-C |
| Homo sapiens | snord96a | GGA-GTG-AGG-ACA-TGT-CCT-GC |
| Homo sapiens | snord95 | GGT-GCT-GAA-ATC-CAG-AGG-CTG-TTT-C |
| Homo sapiens | snord33 | TGA-GAT-GAC-TCT-ACA-TGC-ACT-ACC |
| Homo sapiens | snord66 | CAC-CAT-GAT-GA-ACT-GAG-GAT |
| Homo sapiens | snord44 | TGA-AGG-TCT-TAA-TTA-GCT-CTA-AC |
| Homo sapiens | snord26 | CTG-ATG-GAT-TAG-TGG-AGA-AAA-C |
| Homo sapiens | snord98 | GAA-ATG-CAG-TGT-GGA-ACA-CAA-TG |

**Temperature specific reverse snoRNA primer sequences**

| **Species** | **name** | **sequence** |
| --- | --- | --- |
| **Homo sapiens** | **UNIR 45** | **TAG-TTA-AGC-TTG-GTA-CCG-AG** |
| **Homo sapiens** | **UNIR 50** | **AAT-TCT-AGA-GCT-CGA-GGC-AGG** |
| **Homo sapiens** | **UNIR 55** | **CGA-ATT-CTA-GAG-CTC-GAG-GCA-GG** |
| **Homo sapiens** | **UNIR 60** | **CGA-ATT-CTA-GAG-CTC-GAG-GCA-GGC-GAC** |
| **Homo sapiens** | **UNIR 65** | **CTA-GAG-CTC-GAG-GCA-GGC-GAC-ATG-GCT-GGC** |
| **Homo sapiens** | **RTQ primer** | **CGA-ATT-CTA-GAG-CTC-GAG-GCA-GGC-GAC-ATG-GCT-GGC-TAG-TTA-AGC-TTG-GTA-CCG-AGC-TCG-GAT-CCA-CTA-GTC-CTT-TTT-TTT-TTT-TTT-TTT-TTT-TTT-TTV-N** |

**Overexpression -Coding sequences of human SNORD26 and SNORD96A, flanked by XhoI and MluI restriction sites**

| SNORD26 | CTCGAGCTACGGGGATGATTTTACGAACTGAACTCTCTCTTTCTGATGGATTAGTGGAGAAAACAGAAAATTCTGAGTAGCACGCGT |
| --- | --- |
| SNORD96A | CTCGAGGGTCCTGGTGATGACAGATGGCATTGTCAGCCAATCCCCAAGTGGGAGTGAGGACATGTCCTGCAATTCTGAAGGGATTACGCGT |

**SNORD26 and SNORD96A ASO sequences**

| SNORD26 ASO | 5'- (2'OMeC)(2'OMeC)(2'OMeA) - (2'OMeU)(2'OMeC)A* - G*A*A* - A*G*A* - G*A*G* - (2'OMeA)(2'OMeG)(2'OMeU) - (2'OMeU)(2'OMeC) - 3' |
| --- | --- |
| SNORD26 scrambled | 5'- (2'OMeA)(2'OMeG)(2'OMeG) - (2'OMeA)(2'OMeG)T* - A*G*T* -A*C*G* -A*A*C* - (2'OMeU)(2'OMeA)(2'OMeC) - (2'OMeC)(2'OMeA) - 3' |
| SNORD96A ASO | 5'- (2'OMeA)(2'OMeC)(2'OMeA) - (2'OMeA(2'OMeU)G* - C*C*A* - T*C*T* - G*T*C* - (2'OMeA)(2'OMeU)(2'OMeC) - (2'OMeA)(2'OMeC) - 3' |
| SNORD96A scrambled | 5’-(2’OMeA)*(2’OMeC)*(2’OMeA)*(2’OMeG)*(2’OMeU)*G*T*C*-G*C*G*-T*T*A*-G*(2’OMeA)*(2’OMeU)*(2’OMeA)*(2’OMeA)*(2’OMeU)- 3’ |
