## Supplementary 2 for "SnoRNA signatures in cartilage ageing and osteoarthritis"

**Supplementary File 2- Donor details for human chondrocyte studies**

A. Donors used for rRNA expression studies

| **Status** | **Sex** | **Age (years)** |
| --- | --- | --- |
| Non-OA | male | 45 |
| Non-OA | male | 56 |
| Non-OA | female | 65 |
| Non-OA | female | 69 |
| OA | male | 61 |
| OA | male | 67 |
| OA | female | 68 |
| OA | female | 71 |

B. Donors used for IL-1β and control versus OA synovial fluid studies

| **Status** | **Sex** | **Age (years)** |
| --- | --- | --- |
| Non-OA | male | 45 |
| Non-OA | male | 56 |
| Non-OA | female | 65 |
| Non-OA | female | 69 |

C. Donors used for Non-OA synovial fluid versus OA synovial fluid studies

| **Status** | **Sex** | **Age (years)** |
| --- | --- | --- |
| Non-OA | male | 24 |
| Non-OA | male | 58 |
| Non-OA | female | 23 |
| Non-OA | female | 62 |

D. Donors used for provision of Non-OA and OA synovial fluid

| **Status** | **Sex** | **Age (years)** |
| --- | --- | --- |
| OA | male | 76 |
| OA | male | 62 |
| OA | male | 63 |
| OA | male | 57 |
| OA | female | 50 |
| OA | female | 77 |
| OA | female | 60 |
| OA | female | 73 |
| OA | female | 62 |
| OA | female | 74 |
| Non-OA | male | 62 |
| Non-OA | male | 58 |
| Non-OA | male | 64 |
| Non-OA | male | 59 |
| Non-OA | male | 60 |
| Non-OA | male | 54 |
| Non-OA | male | 51 |
| Non-OA | male | 39 |
| Non-OA | male | 56 |
| Non-OA | male | 64 |

E. Donors used for cultures studies to determine the effect of oxygen, serum on snoRNA gene expression

| **Status** | **Sex** | **Age (years)** |
| --- | --- | --- |
| Non-OA | male | 48 |
| Non-OA | male | 14 |
| Non-OA | female | 13 |

F. Donors use for ASO experiments (all passage 2).

| **Status** | **Sex** | **Age (years)** |
| --- | --- | --- |
| Non-OA | female | 19 |
| Non-OA | female | 26 |
| Non-OA | male | 20 |
| Non-OA | male | 24 |
| Non-OA | male | 23 |
| Non-OA | male | 21 |

G. Donors used for pooled overexpression experiments (all passage 2).

| **Status** | **Sex** | **Age (years)** |
| --- | --- | --- |
| OA | male | 58 |
| OA | male | 78 |
| OA | male | 68 |
| OA | female | 70 |
| OA | female | 73 |
