## Supplementary 3 for "SnoRNA signatures in cartilage ageing and osteoarthritis"

**Supplementary File 3. Details of age, K&L Score, Modified Mankin’s Score and Outerbridge Score for donors used in the microarray**

| **Sample name** | **Age** | **Disease status** | **K&L Score** | **Mankin Score** | | | | **Outerbridge Score** | **Subgroup** |
| --- | --- | --- | --- | --- | --- | --- | --- | --- | --- |
|  |  |  |  | **Scorer 1; 1st** | **Scorer 1; 2nd** | **Scorer 2; 1st** | **Scorer 2; 2nd** |  |  |
| 26U | 52 | U | 4 | N/A | N/A | N/A | N/A | 4 | U1 |
| 28U | 59 | U | 2 | 5 | 4 | 8 | 7 | 3 | U1 |
| 39U | 71 | U | 4 | 3 | 4 | 4 | 3 | 2 | U1 |
| 31U | 73 | U | N/A | N/A | N/A | N/A | N/A | N/A | U2 |
| 38U | 53 | U | 3 | 7 | 7 | 8 | 6 | 2 | U2 |
| 55U | 61 | U | 3 | 2 | 2 | 1 | 1 | 2 | U2 |
| 22U | 58 | U | N/A | N/A | N/A | N/A | N/A | N/A | U3 |
| 41U | 63 | U | 3 | 7 | 7 | 7 | 7 | 1 | U3 |
| 42U | 68 | U | 3 | 9 | 9 | 8 | 9 | 4 | U3 |
| 48U | 68 | U | 2 | 1 | 2 | 2 | 2 | 1 | U3 |
| 22P | 58 | P | N/A | N/A | N/A | N/A | N/A | N/A | P1 |
| 26P | 52 | P | 2 | 4 | 4 | 3 | 3 | 2 | P1 |
| 28P | 59 | P | 1 | 1 | 1 | 2 | 2 | 0 | P1 |
| 38P | 53 | P | 1 | 1 | 2 | 2 | 1 | 1 | P1 |
| 31P | 73 | P | N/A | N/A | N/A | N/A | N/A | N/A | P2 |
| 39P | 71 | P | 1 | 3 | 3 | 4 | 4 | 1 | P2 |
| 41P | 63 | P | 3 | 4 | 4 | 4 | 5 | 0 | P2 |
| 42P | 68 | P | 3 | 5 | 5 | 3 | 5 | 1 | P2 |
| 48P | 68 | P | 2 | 5 | 2 | 4 | 2 | 0 | P2 |
| 55P | 61 | P | 2 | 1 | 1 | 1 | 3 | 1 | P2 |
| 8Y | 28 | Y | N/A | N/A | N/A | N/A | N/A | N/A | Y |
| 9Y | 20 | Y | N/A | N/A | N/A | N/A | N/A | N/A | Y |
| 19Y | 30 | Y | N/A | N/A | N/A | N/A | N/A | N/A | Y |
| 24Y | 25 | Y | N/A | N/A | N/A | N/A | N/A | N/A | Y |
| 25Y | 25 | Y | N/A | N/A | N/A | N/A | N/A | N/A | Y |
| 27Y | 21 | Y | N/A | N/A | N/A | N/A | N/A | N/A | Y |
| 37Y | 18 | Y | N/A | N/A | N/A | N/A | N/A | N/A | Y |
| 56Y | 24 | Y | N/A | N/A | N/A | N/A | N/A | N/A | Y |
| 60Y | 23 | Y | N/A | N/A | N/A | N/A | N/A | N/A | Y |

N/A; no assessment
