## Supplementary 5 for "SnoRNA signatures in cartilage ageing and osteoarthritis"

**Supplementary File 5. SnoRNAs DE in both Y versus P1 and Y versus P2.**

| **Name** | **logFC.Y.VS.P1** | **FDR.Y.VS.P1** | **logFC.Y.VS.P2** | **FDR.Y.VS.P2** | **Class** |
| --- | --- | --- | --- | --- | --- |
| snora48 | 1.8 | 0.01 | 1.3 | 0.03 | HAcaBox |
| snora50 | 0.8 | 0.04 | 0.7 | 0.03 | HAcaBox |
| snora70 | 1.6 | 0.00 | 0.8 | 0.04 | snoRNA |
| snord105B | 3.1 | 0.01 | 1.3 | 0.00 | CDBox |
| snord112 | 1.1 | 0.01 | 2.6 | 0.01 | CDBox |
| snord113_4 | 1.5 | 0.01 | 0.9 | 0.01 | CDBox |
| snord113_7 | 1.9 | 0.01 | 1.4 | 0.01 | CDBox |
| snord113_8 | 1.2 | 0.02 | 1.8 | 0.01 | CDBox |
| snord113_9 | 1.2 | 0.02 | 1.0 | 0.02 | CDBox |
| snord114_12 | 2.4 | 0.00 | 0.9 | 0.03 | CDBox |
| snord114_14 | 5.9 | 0.00 | 1.8 | 0.01 | CDBox |
| snord114_17 | 1.6 | 0.00 | 2.9 | 0.00 | CDBox |
| snord114_21 | 3.0 | 0.00 | 1.3 | 0.00 | CDBox |
| snord114_22 | 3.2 | 0.00 | 3.0 | 0.00 | CDBox |
| snord114_24 | -1.1 | 0.02 | 3.1 | 0.00 | CDBox |
| snord114_26 | 2.2 | 0.00 | -1.1 | 0.01 | CDBox |
| snord114_28 | 2.6 | 0.00 | 1.6 | 0.01 | CDBox |
| snord114_3 | 1.7 | 0.01 | 2.8 | 0.00 | CDBox |
| snord1B | 0.9 | 0.01 | 1.8 | 0.00 | CDBox |
| snord36 | 5.7 | 0.00 | 0.7 | 0.04 | snoRNA |
| snord36A | 1.3 | 0.03 | 1.3 | 0.01 | CDBox |
| snord38B | 4.8 | 0.00 | 1.3 | 0.01 | CDBox |
| snord45 | -0.6 | 0.00 | 4.0 | 0.00 | snoRNA |
| snord49B | 1.7 | 0.00 | -0.5 | 0.01 | CDBox |
| snord56 | 4.0 | 0.00 | 1.3 | 0.00 | CDBox |
| snord59A | 5.9 | 0.00 | 2.2 | 0.03 | CDBox |
| snord62 | 1.6 | 0.03 | 3.1 | 0.02 | snoRNA |
| snord63 | 5.0 | 0.00 | 3.2 | 0.04 | CDBox |
| snord75 | 3.8 | 0.00 | 2.5 | 0.04 | CDBox |
| snord82 | 2.0 | 0.00 | 1.4 | 0.01 | CDBox |
| snord83 | 4.7 | 0.00 | 3.9 | 0.00 | snoRNA |
| snord98 | 2.6 | 0.00 | 2.2 | 0.01 | CDBox |
| snoU13 | -4.4 | 0.00 | -2.0 | 0.03 | snoRNA |

Abbreviations- Y; young normal, P; protected (old normal) LFC; log fold change, FDR; false discovery rate
