## Supplementary 6 for "SnoRNA signatures in cartilage ageing and osteoarthritis"

**Supplementary File 6** Topographical snoRNA gene expression in non-OA cartilage. RNA extracted from equine metacarpophalangeal joints; low load area was lateral condylar area and high load area was medial condylar area n=5. SnoRNA gene expression relative to U6 and protein coding genes to GAPDH. Data represents mean ± standard error of mean. Statistical analyses undertaken following normality testing with a Mann Whitney Test.

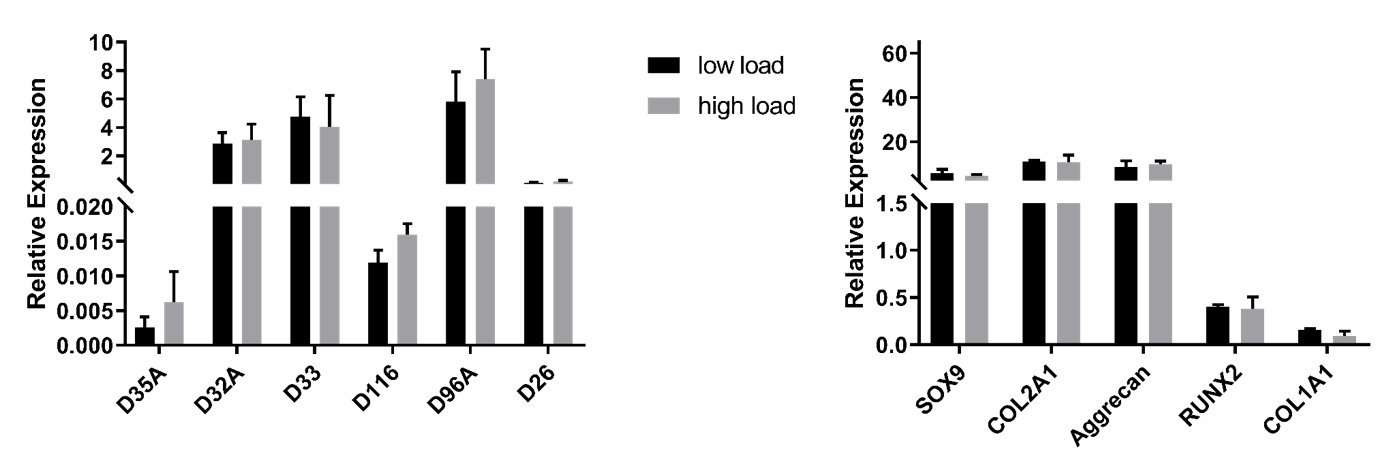
