## Supplementary 7 for "SnoRNA signatures in cartilage ageing and osteoarthritis"

Supplementary File 7. Expression of mRNA for indicated genes following A. IL-1β treatment or B. OA synovial fluid treatment measured using qRT-PCR. Treatment of non-OA HAC with IL-1β (10ng/ml) for 24 hours or with 20% OA synovial fluid (SF) (derived from a pool of ten donors) for 24 hours. Gene expression changes were measured using 2^-ΔCT expression relative to cyclophilin. Data represents the mean + standard error mean, P values indicated as follows; p<0.05 *, p<0.01 **. Within the graphs black represents control and grey treatments.

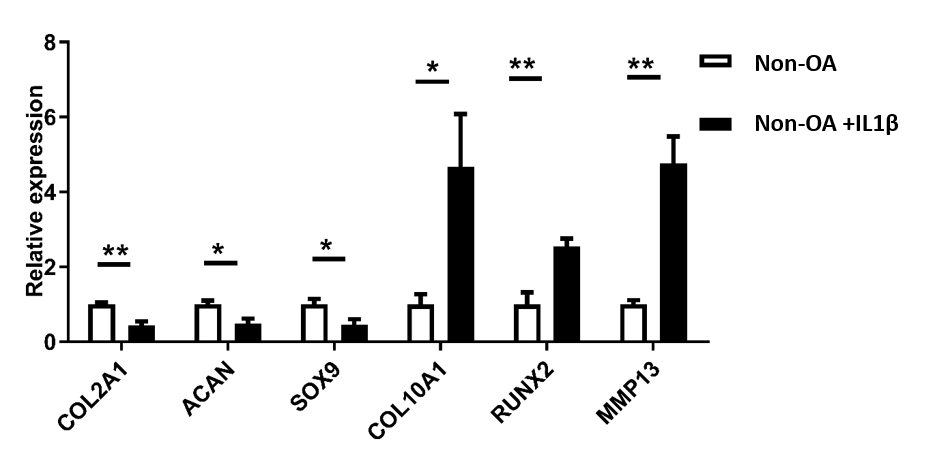

B.

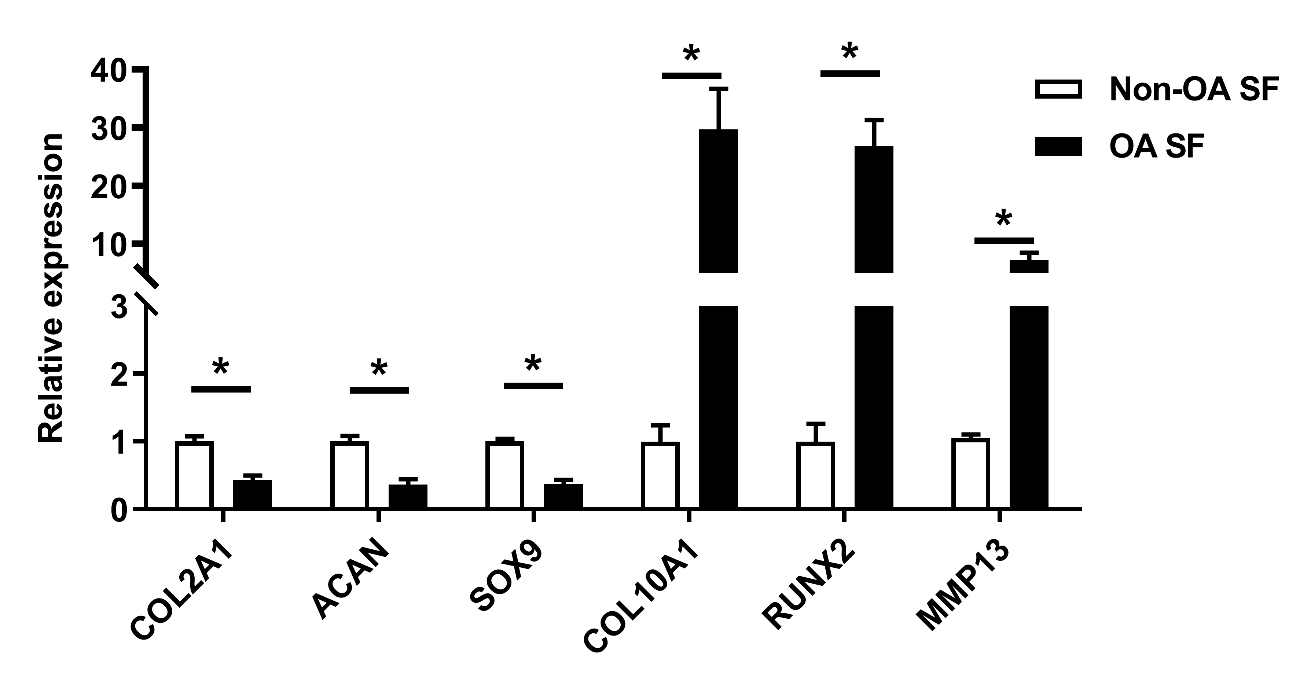
