## Supplementary 8 for "SnoRNA signatures in cartilage ageing and osteoarthritis"

**Supplementary File 8**. Effect of chondrocyte passage on snoRNA gene expression. There was little effect of passage on snoRNA expression. Passaged chondrocyte (n=5) snoRNA gene expression was relative to U6 and protein coding genes to GAPDH. Histogram represents mean ± standard error of mean. Statistical analyses undertaken using ANOVA.

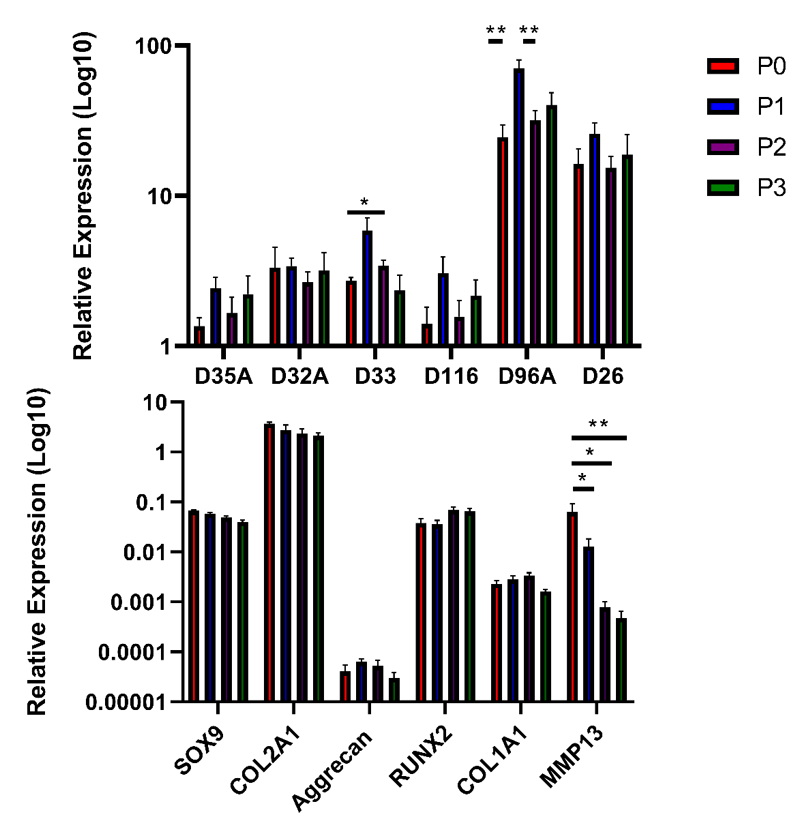
