## Supplementary 9 for "SnoRNA signatures in cartilage ageing and osteoarthritis"

Supplementary File 9. The effect of oxygen tension and serum in cell culture had no effect on selected snoRNA gene expression of non-OA HAC. A. P1 chondrocytes (n=3) were subject to 5% (low oxygen) and 20% (high oxygen) with (plus serum) and without (serum-free) 10% FCS. SnoRNA gene expression was relative to U6 and protein coding genes to GAPDH. Histogram represents mean ± standard error of mean. B. Represents assessment of chondrogenic and hypertrophic gene expression in the same samples. Statistical analyses undertaken using a one-way ANOVA.

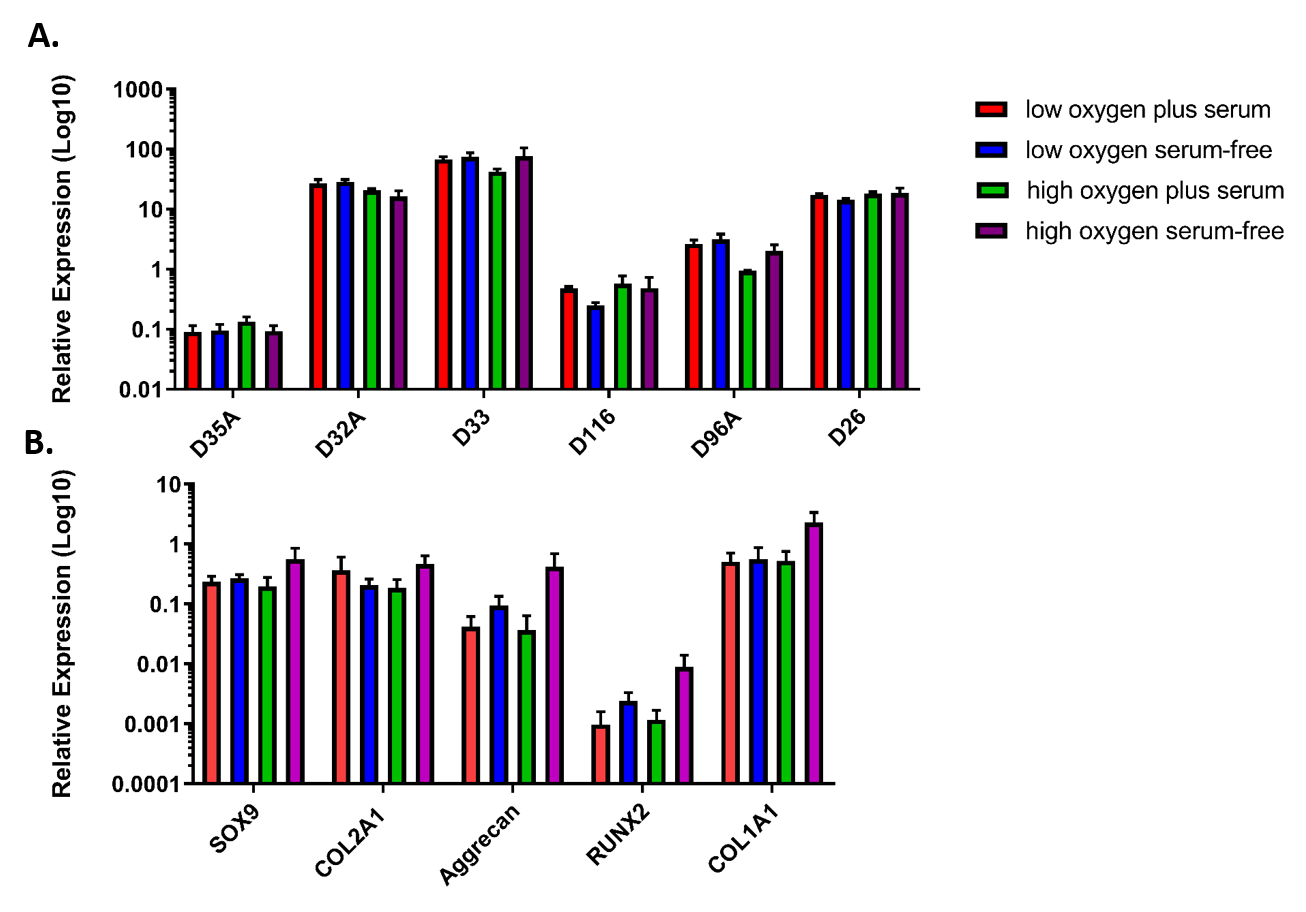
